# Insights into the Molecular Mechanism of Pulmonary Vein Stenosis in Pediatric Patients

**DOI:** 10.1101/2024.10.09.617404

**Authors:** Alyssa B Kalustian, Pengfei Ji, Lalita Wadhwa, Christopher A. Caldarone, Manish Bansal, Athar M. Qureshi, Jeffrey S. Heinle, Ravi K. Birla

## Abstract

**Background:** Pediatric pulmonary vein stenosis (PVS) is associated with substantial morbidity and mortality for the subset of patients with recurrent or progressive disease. The molecular mechanisms underlying the development and trajectory of PVS remain unclear. This study characterizes the transcriptome of clinical and phenotypic subtypes of PVS.

**Methods:** Bulk RNA sequencing analysis was performed on human pulmonary vein tissue samples obtained from surgical interventions for pediatric patients with PVS. Transcriptomic profiles were compared for primary versus post-repair PVS as well as aggressive versus non-aggressive clinical phenotypes. Principal component analysis was performed, the differential gene expression quantified, and pathway analysis conducted based on gene ontology, KEGG and Reactome.

**Results:** When comparing AggPPVS vs NonAggPPVS, differences were noted in the genes associated in extracellular matrix regulation and PIEZO1, a mechanosensitive receptor present in endothelial cells. In addition, there were notable changes in cardiac muscle contractility, calcium handling, respiratory and energy metabolism. These results point to a potential mechanism for aggressive PPVS phenotype, due to an overexpression of PIEOZ1 in response to elevated shear stress, subsequent activation of intracellular signaling pathways and leading to reduced contractility and intracellular calcium transients with cardiomyocytes.

**Conclusions:** The result of this study suggests that aggressive PPVS phenotype is caused by an increase in PIEZO1 expression and subsequent changes in extracellular matrix production, heart muscle contractility and changes in calcium transients within cardiomyocytes. These results provide a potential target for therapeutic invention for primary PPVS by inhibiting the activity of PIEZO1. This could potentially reduce morbidity in this patient population.

## Introduction

Pediatric pulmonary vein stenosis (PVS) is a rare, poorly understood disease process associated with high recurrence, morbidity, and mortality^1^. Obstructive neo-intimal lesions in the pulmonary veins can lead to pulmonary hypertension and right heart failure, necessitating intensive surveillance, multimodal therapy, and multiple interventions^1^. There are two overarching categories of PVS, each differing somewhat in their hypothesized pathogenesis^1^. Primary or congenital PVS is characterized by luminal narrowing that develops in the absence of prior surgical intervention or instrumentation, with a potentially distinct subgroup associated with prematurity and other pulmonary structural anomalies^2^. Abnormal or disrupted embryonic signaling is thought to underly primary PVS^2^. By contrast, post-repair PVS (PRPVS) develops following surgical intervention on the pulmonary veins, including 5-15% of patients with traditionally repaired total anomalous pulmonary venous connection (TAPVC)^3^.

Mechanical factors such as turbulent flow patterns^4^ and abnormal proliferative/fibrotic response^5^ to suture foreign bodies likely drive this process. Implementation of sutureless technique for TAPVC repair has been associated with reduced incidence of PRPVS, supporting this hypothesis^6^. Within both primary and PRPVS, a subset of patients has an aggressive, treatment-refractory phenotype characterized by disease progression (in severity or extent) and recurrence despite interventions.

While several clinical risk-factors have been identified^7^, no biomarkers have been associated with disease prognosis or outcomes. Genetic syndromes and chromosomal abnormalities are not uncommon in patients with PVS^8^, and a locus of interest was identified in several related patients with PVS and lymphatic anomalies^9^; however, no other genetic mutations have been reliably implicated in PVS. Histologically, the lesions are marked by myofibroblast-like cells and extracellular matrix deposition. Possible cellular pathways involved in pathogenesis include activation of tyrosine kinase receptors^10^, activation of mammalian target of rapamycin^11^, EndoMT^12^ and TGF-beta signaling^5^.

The delineation into subtypes, including primary and post-repair have not been clearly defined, beyond the clinical observations that primary PVS is congenital, present at the time of birth while post-repair PVS is acquired in response to surgical intervention. While aggressive phenotype is observed in both primary and post-repair PVS, the commonalities or differences between aggressive and nonaggressive phenotype in each of these cases is not known. Such information is important to understand and manage PVS and develop effective treatment strategies.

An understanding of the changes of in the gene expression of PVS patients is a necessary prerequisite to the management and treatment this patient population. This study aims to characterize the transcriptome of PVS to identify potential biomarkers of disease phenotype and clues to disease pathogenesis. To accomplish this, we correlated bulk RNA sequencing analysis of human pulmonary vein specimens obtained during surgical intervention with clinical outcomes. The bulk-RNA sequencing and subsequent analysis were used to identify specific genes and pathways that are differentially regulated between PPVS and PRPVS and the aggressive and nonaggressive phenotype in both cases. The results of this study provide important insight into the potential molecular signaling pathways and support the development of potential therapies for the treatment of PVS.

The specific objective of this study is to use bulk-RNA sequencing of pulmonary vein samples from PVS patients and identify differences in the gene profile based on primary vs post-repair and aggressive vs nonaggressive phenotype.

## Methods

### Study Design

Following approval by Institutional Review Board (protocol H-52781, approved 12/9/2022), pulmonary vein tissue was collected from the Texas Children’s Hospital Heart Center Biorepository. The pulmonary tissue was excised at the boundary with the left atrium and also contained atrial tissue. Written informed consent for the collection, preservation, long-term storage, and future research use of surgical biospecimens was obtained from the patient’s parent or legal guardian prior to surgery, and verbal assent was obtained from children older than 6 years of age (Institutional Review Board Protocol H-26502). In total, 6 pulmonary vein tissue samples obtained from 6 unique patients were identified and 1 control from a donor heart.

### Clinical Data and Definitions

Pertinent demographic, clinical, interventional, and follow-up information was abstracted from medical records of the patients with eligible samples. This information was used to determine clinical subgroup, phenotype, and outcomes. Clinical subtypes were classified as PPVS or PRPVS based on history of prior surgical intervention for TAPVC or PAPVC at time of PVS diagnosis (**Figure 1A**). Aggressive phenotype was defined as having any of the following criteria based on retrospective review through last follow-up (**Figure 1B**): 2 or more catheter or operative interventions for PVS within any one-year interval, treatment-refractory pulmonary hypertension (persistent despite multiple agents), progression of venous disease from focal/proximal to diffuse/distal or complete occlusion/atresia, progression from single- to multi-vein disease, or progression from unilateral to bilateral disease. Of the 6 patients with pulmonary vein tissue samples, 4 had primary PVS and 2 had PRPVS.

**Figure 1:**
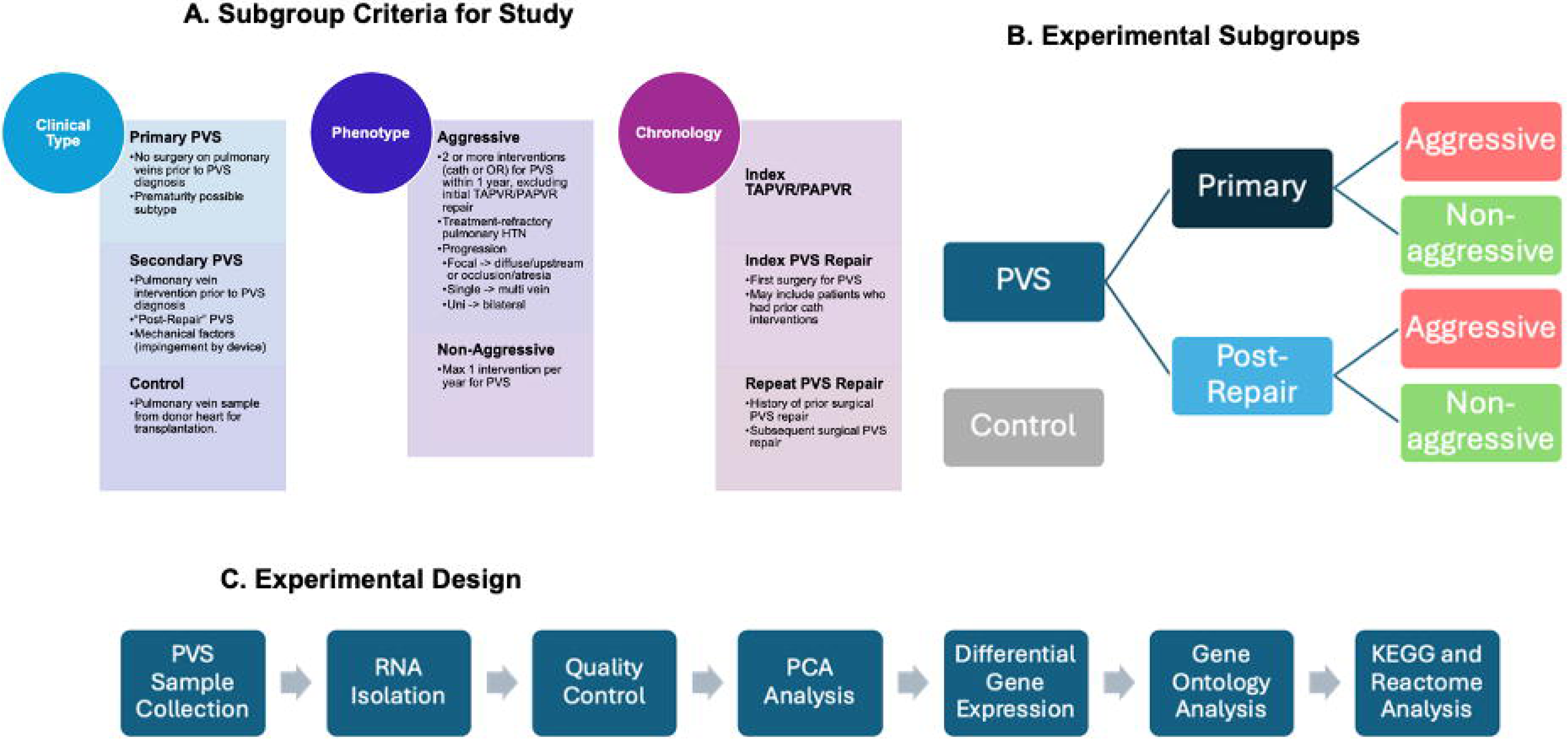
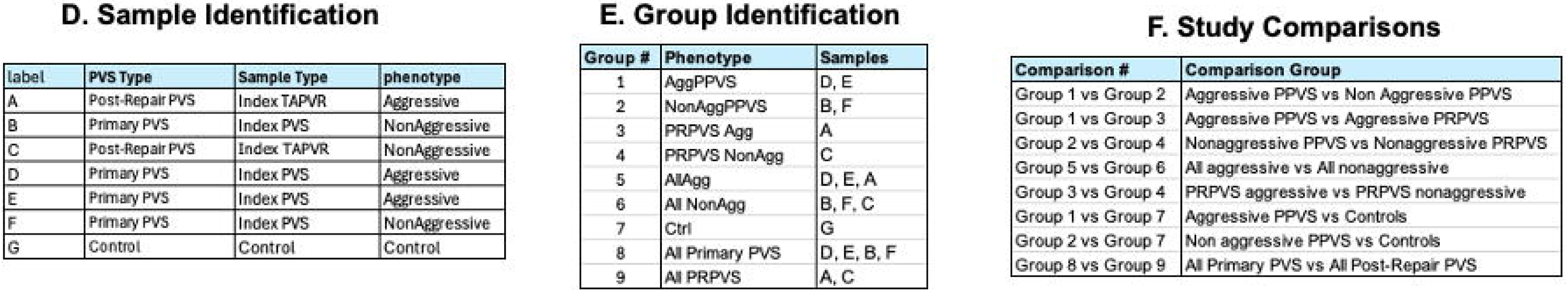
Study Design: (A) Subgroup Criteria for Study. The table shows the clinical type, phenotype and chronology. **(B) Experimental Subgroups:** We included primary and post-repair PVS and for both groups, had samples for both aggressive and nonaggressive phenotype. **(C) Experimental Design:** PVS samples were collected from the biobank and RNA isolation conducted and checked for quality control. Bulk-RNA sequencing was performed, and the analysis consisted of PCA analysis, differential gene expression, gene ontology analysis and KEGG and Reactome analysis. **(D) Sample Identification:** 7 samples were used for this study and labelled as A-G and categorized as primary PVS, post-repair PVS and further categorized as aggressive or nonaggressive. **(E) Group Identification:** 9 groups were identified, as listed in the table and the respective samples for each group are identified in the table as well. ^29^ **Study Comparisons:** 8 comparisons were studies and identified in the Table and the specific comparison groups are also identified in the table.

### Experimental Design

As shown in **Figure 1C**, once all PVS samples were recovered from the biobank, RNA extraction was performed, followed by quality control. Bulk-RNA sequencing was conducted, and the results used for PCA analysis, differential gene expression, gene ontology analysis and KEGG and Reactome analysis.

### Extraction of RNA from Tissue Specimens

1 mL of TRIzol™ reagent was added per 50–100 mg of tissue to the sample and the sample homogenized using a homogenizer. The homogenized sample was incubated for 5 minutes to allow complete dissociation of the nucleoproteins complex. After this, 0.2 mL of chloroform was added per 1 mL of TRIzol™ reagent used for lysis, the cap of the conical tube secured and then the solution was thoroughly mixed by manually shaking for 2-3 minutes. Next, the sample was centrifuged for 15 minutes at 12,000g at 4°C. The process resulted in separation of the tissue suspension into a lower phenol-chloroform, an interphase layer in the middle and a colorless upper aqueous phase. The upper aqueous phase was carefully aspirated and transferred to a clean conical and 0.5 ml of isopropanol added per 1 ml of TRIzol^TM^ reagent, and this new tissue suspension incubated for 10 minutes at 4°C. This tissue suspension was centrifuged for 10 minutes at 12,000g at 4°C, resuspended in 1 ml of 75% ethanol per 1 ml of TRIzol^TM^ reagent. Finally, the supernatant was discarded, and the pellet resuspended in 50 ul of RNase-free water, with 0.1 mM EDTA and 0.5% SDS solution by pipetting up and down multiple times. RNA samples were quantified by absorbance without prior dilution using the NanoDrop™ Spectrophotometer. The ratio of A260/A280 within the range of 1.8-2.0 and A230/280 <1.6 were used for this study.

### Bulk RNA Sequencing

Purified RNA samples were shipped to Novogene Inc on dry ice for bulk RNA sequencing. Data analysis was also conducted by Novogene, and plots were generated for principal component analysis, volcano plots, heatmaps, gene ontology analysis, KEGG pathway analysis and Reactome pathway analysis.

### Statistical Analysis

Statistical analyses were conducted using R (R Foundation for Statistical Computing, Vienna, Austria) // Microsoft Excel (Microsoft Corp, Redmond, Washington) // STATA version *** software (StataCorp LLC, College Station, Texas). Continuous and categorical variables are reported as median and total range or absolute and relative frequencies, respectively. (Comparative methods). A p-value < 0.05 was considered statistically significant.

## Results

### Clinical Course

Pertinent clinical details, including comorbidities and PVS trajectory are summarized for the 6 included patients in **Table 1**. Sample identification, group identification and study comparisons are presented in **Figure 1D-F**. Four patients were male. All 4 patients with primary PVS (#1-4) had additional congenital cardiac lesions as well as genetic syndromes, including VACTERL (n=2), Turner Syndrome (n=1), and Pierre-Robin Syndrome (n=1).

**Table 1:** Clinical Details.

| **Patient** | **PVS Type** | **Phenotype** | **Comorbidities** | **PV Interventions Prior to Sample Collection** | **Operations at Sample Collection** | **Age at sample collection (months)** | **# PVS Interventions Post-Sample Collection** |
| --- | --- | --- | --- | --- | --- | --- | --- |
| 1 | Primary | Aggressive | Turner Syndrome, PAPVC (LUPV), coarctation | Balloon angioplasty right sided veins | Index PVS repair, fenestrated ASD closure | 6.9 | 3 |
| 2 | Primary | Non-aggressive | VACTERL, ASD | None | Index PVS repair, fenestrated ASD closure | 12.4 | 0 |
| 3 | Primary | Aggressive | Pierre-Robin, Diamond-Blackfan anemia, ASD, interrupted IVC, dysplastic TV | None | Index PVS repair, fenestrated ASD closure | 1.1 | 2 |
| 4 | Primary | Non-aggressive | VACTERL, TV atresia, PS, VSD, PDA | None | Index PVS repair, Glenn, PA plasty, atrial septectomy | 9.8 | 1 |
| 5 | Post-Repair | Non-aggressive | Mixed TAPVC | TAPVC repair | Index repair of PR-PVS, completion PAPVC repair | 34.9 | 0 |
| 6 | Post-Repair | Aggressive | Cardiac TAPVC | (1) TAPVC repair (coronary sinus unroofing)<br>(2) Balloon angioplasty stenotic venous confluence<br>(3) Reoperation for confluence stenosis<br>(4) Stent placement | Redo repair of PR-PVS, reconstruction atrial septum | 11.8 | 17 |

\*Unrepaired single-vein PAPVC to innominate vein, this vein not affected by PVS
Abbreviations: ASD = atrial septal defect; PA = pulmonary artery; PDA = patent ductus arteriosus; PS = pulmonary stenosis; TV = tricuspid valve; VACTERL = vertebral defects, anal atresia, cardiac defects, tracheo-esophageal fistula, renal anomalies, and limb abnormalities; VSD = ventricular septal defect

At time of sample collection, all 6 patients had multi-vein, bilateral involvement of their PVS. Median age at sample operation was 10.8 months (range 1.1 month – 2.9 years). Tissue samples were collected from the index operative PVS repair for all 4 primary PVS patients, though Patient 1 had a transcatheter balloon angioplasty prior to surgery. Of the 2 patients with PR-PVS (#5-6), patient 5 underwent early neonatal partial repair of mixed obstructed TAPVC (infradiaphragmatic confluence of right upper/lower and left lower pulmonary veins relocated to left atrium, left upper pulmonary vein left draining via ascending vertical vein to the innominate) without intervening interventions prior to sample collection at operative resection of confluence ostial stenosis and completion TAPVC repair at age 3 years. Patient 6 had initial repair of TAPVC to the coronary sinus around 1 month of age followed by multiple interventions for PRPVS (multiple transcatheter and one reoperation) prior to sample collection at an operation to resect prominent ostial fibrotic stenosis of the venous confluence and perform sutureless repair for left lower pulmonary vein atresia.

Median follow-up time from sample collection was 4 years (range 1.4 - 8.5 years) for survivors. One patient (#3) with numerous severe extracardiac comorbidities died approximately 4 months postoperatively during their index hospitalization due to progressive primary PVS with refractory pulmonary hypertension which culminated in right ventricular failure and cardiac arrest, despite several transcatheter interventions. Their course was also complicated by infectious complications, including disseminated fungemia. All 3 patients with aggressive PVS (#1, 3, 6) had at multiple PVS reinterventions postoperatively (all transcatheter), with time to first reintervention ranging from 27 days to 3.3 months postoperatively. At latest follow-up, 2 patients (#1, 6) were on sirolimus therapy for aggressive PVS. None of the patients with non-aggressive phenotype (#2, 4, 5) required subsequent PVS intervention.

### Association Between Sample Groups

Principal component analysis (PCA) was used to quantify the association between samples that belong to the same groups; the closer the samples from a single group are to each other and to a single quadrant, the better it is. PCA analysis provides initial evidence of the similarity and/or separation of the samples for each group within the study. There are two key areas for analysis. First, samples within each group should be closely placed, thereby suggesting minimal variability within the group. Second, the variability between groups is assessed based on separation of the samples within each group. The PCA analysis for all 8 comparisons is presented in **Figure 2** and each is described next.

**Figure 2:**
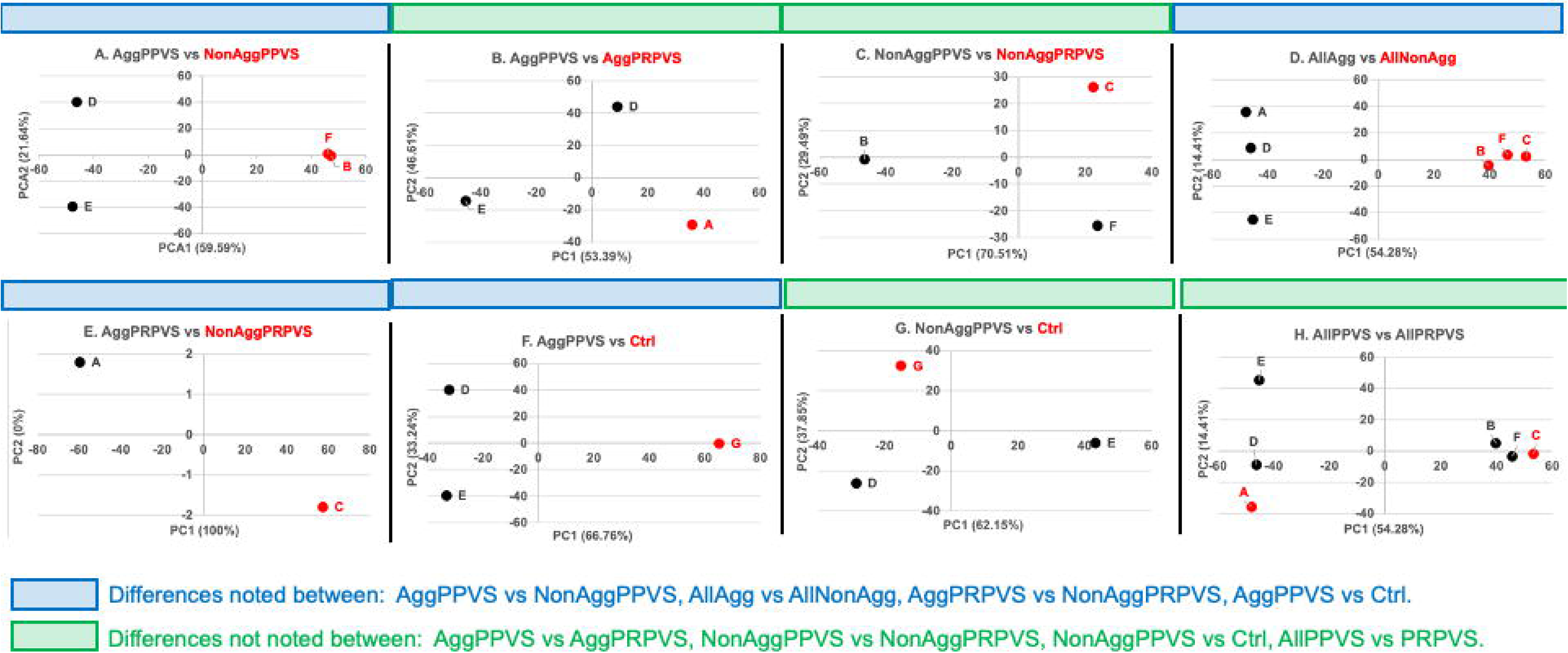
Principal Component Analysis: PCA plots are shown for the following comparisons: (A) AggPPVS vs NonAggPPVS. (B) AggPPVS vs AggPRPVS. (C) NonAggPPVS vs NonAggPRPVS. (D) AllAgg vs AllNonAgg. (E) AggPRPVS vs NonAggPVPVS. AggPPVSvsCtrl. NonAggPPVSvsCtrl. (H) AllPPVS vs AllPRPVS.

**AggPPVS vs NonAggPPVS:** The two samples for AggPPVS (D&E) and the two samples for NonAggPPVS (F&B) are clustered together, thereby suggesting sample uniformity within these two groups. Furthermore, the samples for AggPPVS (D&E) and NonAggPPVS (F&B) were placed on opposite ends of the PCA plots, thereby demonstrating clear separation between these two groups. These results suggest differences between the phenotype of AggPPVS relative to NonAggPPVS. **AggPPVS vs AggPRPVS:** The samples for AggPPVS (D&E) were not placed in proximity with each other when compared with AggPRPVS (A) thereby suggesting that these two groups are similar. This would suggest aggressive phenotype has a common origin in both primary and post-repair PVS. **NonAggPPVS vs NonAggPRPVS:** These results mirrored those for AggPPVS vs AggPRPVS. The samples for NonAggPPVS (F&B) were separated from each other and from the sample for AggPRPVS (C) suggesting similarities between these two groups. Collectively, PCA analysis of AggPPVS vs AggPRPVS and NonAggPPVS vs NonAggPRPVS demonstrate similarities between PPVS and PRPVS, in both cases of a primary or post-repair PVS.

**AllAgg vs AllNonAgg:** The samples for AllAgg (A, D&E) all clustered together and the same was noted for the samples for AllNonAgg (B, C&F), demonstrating agreement within the group. The samples for both groups separated on the PCA plot, placed on either end of the plot. These results demonstrate differences between aggressive vs nonaggressive phenotype in mediating PVS phenotype, irrespective of PPVS or PRPVS. **AggPRPVS vs NonAggPRPVS:** The sample for AggPRPVS (A) and NonAggPRPVS (C) were placed on opposite ends of the PCA plot, suggesting significant differences between these two phenotypes. **AggPPVS vs Ctrl:** The samples for AggPPVS (D&E) were placed in close proximity, suggesting agreement within this group. Furthermore, there was a clear separation between samples for AggPPVS (D&E) relative to the ctrl, suggesting differences between the two groups. **NonAggPPVS vs Ctrl:** The samples for NonAggPPVS (B&F) were similar from the ctrl sample. In the previous comparison, we determined that aggressive PPVS has clear differences relative to controls, whereas the results here demonstrate similarities between nonaggressive PPVS and controls.

**AllPPVS vs AllPRPVS:** Based on our current understanding of PVS, PPVS and PRPVS are believed to be distinct^13^, the former being related to developmental abnormalities while the latter related to injury resulting from surgical interventions. However, our results show that PPVS and PRPVS are similar, more than originally believed. Based on the PCA plot, samples for AllPPVS (D, E, B&F) and AllPRPVS (A&C) show differences.

The results of the PCA analysis show clear differences between 4 groups: AggPPVS vs NonAggPPVS, AllAgg vs AllNonAgg, AggPRPVS vs NonAggPRPVS, AggPPVS vs Ctrl. These results are consistent with on our current understanding of the disease and provide additional evidence supporting clinical observations. The results of the PCA analysis show similarities between 4 groups: AggPPVS vs AggPRPVS, NonAggPPVS vs NonAggPRPVS, NonAggPRPVS vs Ctrl and AllPPVS vs AllPRPVS. These results provide new insight into the similarities between PPVS relative to PRPVS, currently thought to be different based clinical observations.

### Additional Validation of Separation of Experimental Groups

Once we established significant differences between study groups based on PCA analysis, our next objective was to quantify the number of differentially expressed genes (DEGs) between the group tested earlier. We used volcano plots to visualize the number of DEGs and listed the number of upregulated and downregulated genes on these plots. The first step in the analysis of the bulk RNA sequencing data is an in-depth analysis of the total number of individual genes that are differentially expressed. This dataset is used to provide a global overview of the changes in the gene expression pattern and provides a pathway moving forward. The results are presented in **Figure 3** and described here. **AggPPVS vs NonAggPPVS:** there were 2097 upregulated genes and 3292 downregulated genes. **AggPPVS vs AggPRPVS:** there were 237 upregulated genes and 115 downregulated genes. **NonAggPPVS vs NonAggPRPVS:** there were 173 upregulated genes and 20 downregulated genes. **AllAgg vs AllNonAgg:** there were 3529 upregulated genes and 4449 downregulated genes. **AggPRPVS vs NonAggPRPVS:** there were 4445 upregulated genes and 5815 downregulated genes. **AggPPVS vs Ctrl:** there were 1099 upregulated genes and 2352 downregulated genes. **NonAggPPVS vs Ctrl:** there were 147 upregulated genes and 281 downregulated genes. **AllPPVS vs AllPRPVS:** there were 217 upregulated genes and 79 downregulated genes.

**Figure 3:**
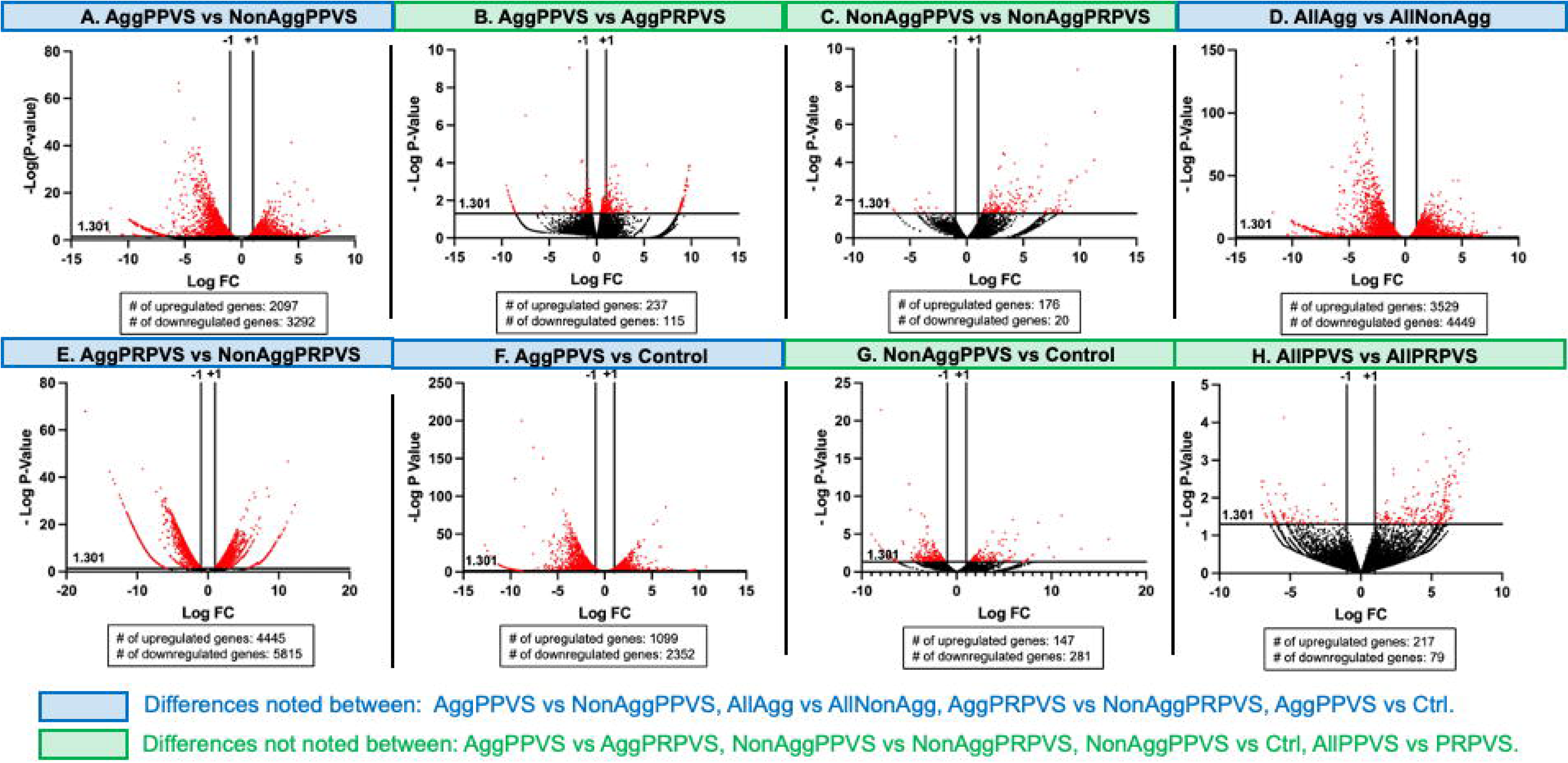
Differentially Expressed Genes: Volcano plots are shown for the following comparisons: (A) AggPPVS vs NonAggPPVS. (B) AggPPVS vs AggPRPVS. (C) NonAggPPVS vs NonAggPRPVS. (D) AllAgg vs AllNonAgg. (E) AggPRPVS vs NonAggPVPVS. AggPPVS vs Ctrl. NonAggPPVS vs Ctrl. (H) AllPPVS vs AllPRPVS.

The results of the DEGs show significant differences between 4 groups, in which case the number of DEGs is in the order of magnitude of thousands: AggPPVS vs NonAggPPVS, AllAgg vs AllNonAgg, AggPRPVS vs NonAggPRPVS, AggPPVS vs Ctrl. The results of the DEGs show significant similarities between 4 groups, in which case the number of DEGs is small, less than 100: AggPPVS vs AggPRPVS, NonAggPPVS vs NonAggPRPVS, NonAggPRPVS vs Ctrl and AllPPVS vs AllPRPVS.

### Funneling Strategy for Further Analysis

We started our analysis based on 8 groups and conducted PCA and DEG analysis to identify the groups and comparisons with the most significant changes. Based on this analysis, we identified 4 groups for further analysis which consisted of: AggPPVS vs NonAggPPVS, AllAgg vs AllNonAgg, AggPRPVS vs NonAggPRPVS, AggPPVS vs Ctrl. These are the only groups that were retained for further analysis.

### Genes Associated with Specific Phenotype

Our objective was to identify the 50 most regulated genes, the 50 most downregulated genes and the 50 most significant genes for our selected groups. The results are presented as a heatmap in **Figure 4**. Each of our 4 selected comparisons are presented independently.

**Figure 4:**
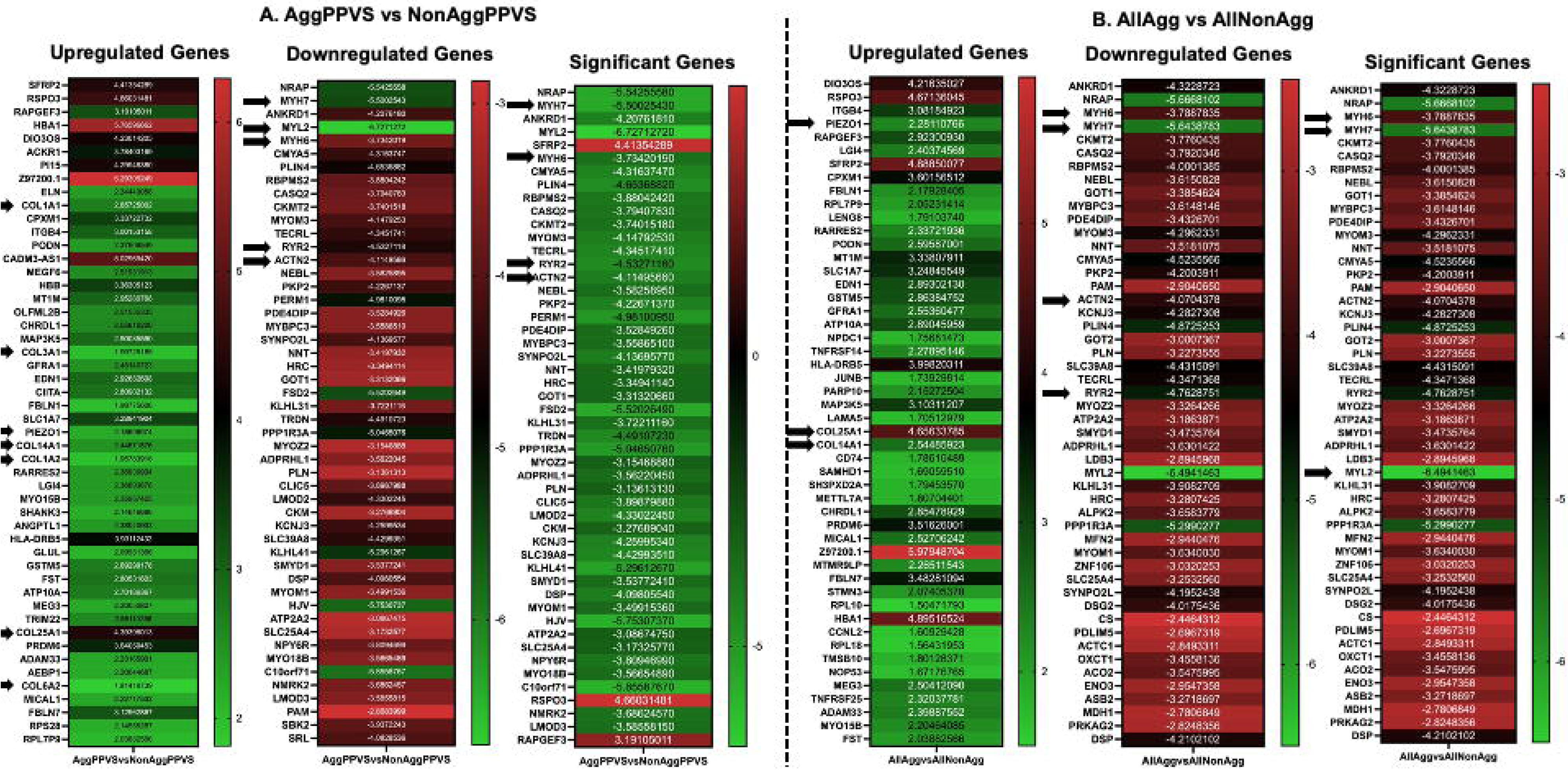
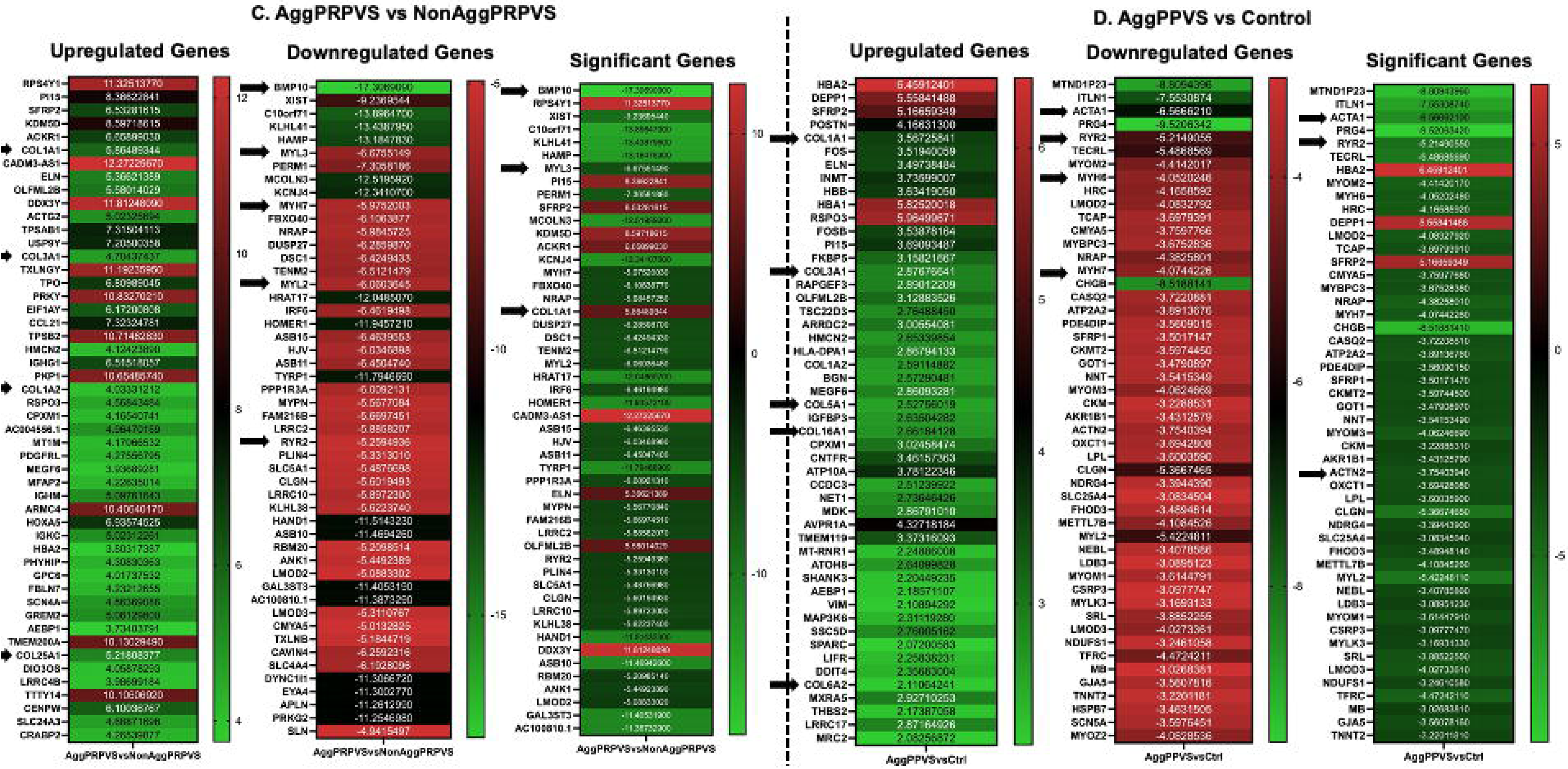
Heatmaps Showing Top 50 Genes: The top 50 upregulated, downregulated and most significant genes are presented for the 4 groups: (A) AggPPVS vs NonAggPPVS. (B) AllAgg vs AllNonAgg. (C) AggPRPVS vs NonAggPRPVS. (D) AggPPVS vs Control.

**AggPPVS vs NonAggPPVS:** Many of the upregulated genes were involved in ECM remodeling and were part of the collagen family and included COL1A1, COL14A1, COL1A2, COL25A1 and COL6A2. Another interesting upregulate gene was PIEZO1, a mechanosensory receptor in endothelial cells. Many of the down regulated genes were also included involved in ECM remodeling and included MYH7, MYL2, MYH6 and ACTN2. Genes associated with suppressed cardiac function was also noted based on the expression pattern of RYR2, which regulates intracellular calcium in cardiomyocytes. This trend was also observed when looking at the most significant genes, as they belonged to either upregulated or downregulated genes. **AllAgg vs AllNonAgg:** The results for this comparison were similar to those for AggPPVSvsNonAggPPVS where ECM proteins and PIEZO1 were upregulated along with many genes associated with ECM remodeling, including segments of collagen COL251A and COL141A. As was the case before, many genes associated with ECM remodeling were also downregulated and included MYH6, MYH7 and ACTN2. In addition, RYR2 was downregulated and this suggested changes in intracellular calcium transients in cardiomyocytes.

**AggPRPVS vs NonAggPRPVS and AggPPVS vs Ctrl:** The trend continued for these two comparisons, with changes in ECM proteins and RYR2 expression. While PIEZO1 expression was upregulated in AggPPVSvsNonAggPPVS and AllAggvsAllNonAgg, it was not identified in AggPRPVSvsNonAggPRPVS and AggPPVSvsCtrl.

### Gene Ontology Analysis

The next step in our data analysis was pathway analysis, looking at variations in individual genes and how these come together to alter specific pathways within cells and subsequently tissue. Gene ontology analysis represents a high-level representation of changes in specific pathways, altered due to changes in the expression of specific genes. GO analysis is divided into three sub-domains, molecular function (MF), cellular component (CC) and biological process (BP). MF represents activities resulting from changes in the specific pathways. CC represents the location within the cellular structures where the MFs are performed, for example, cytoplasm, mitochondria, golgi apparatus etc. BP refers to the specific process that is altered and examples include DNA repair or signal transduction. The results of the GO analysis are presented in **Figure 5**.

**Figure 5:**
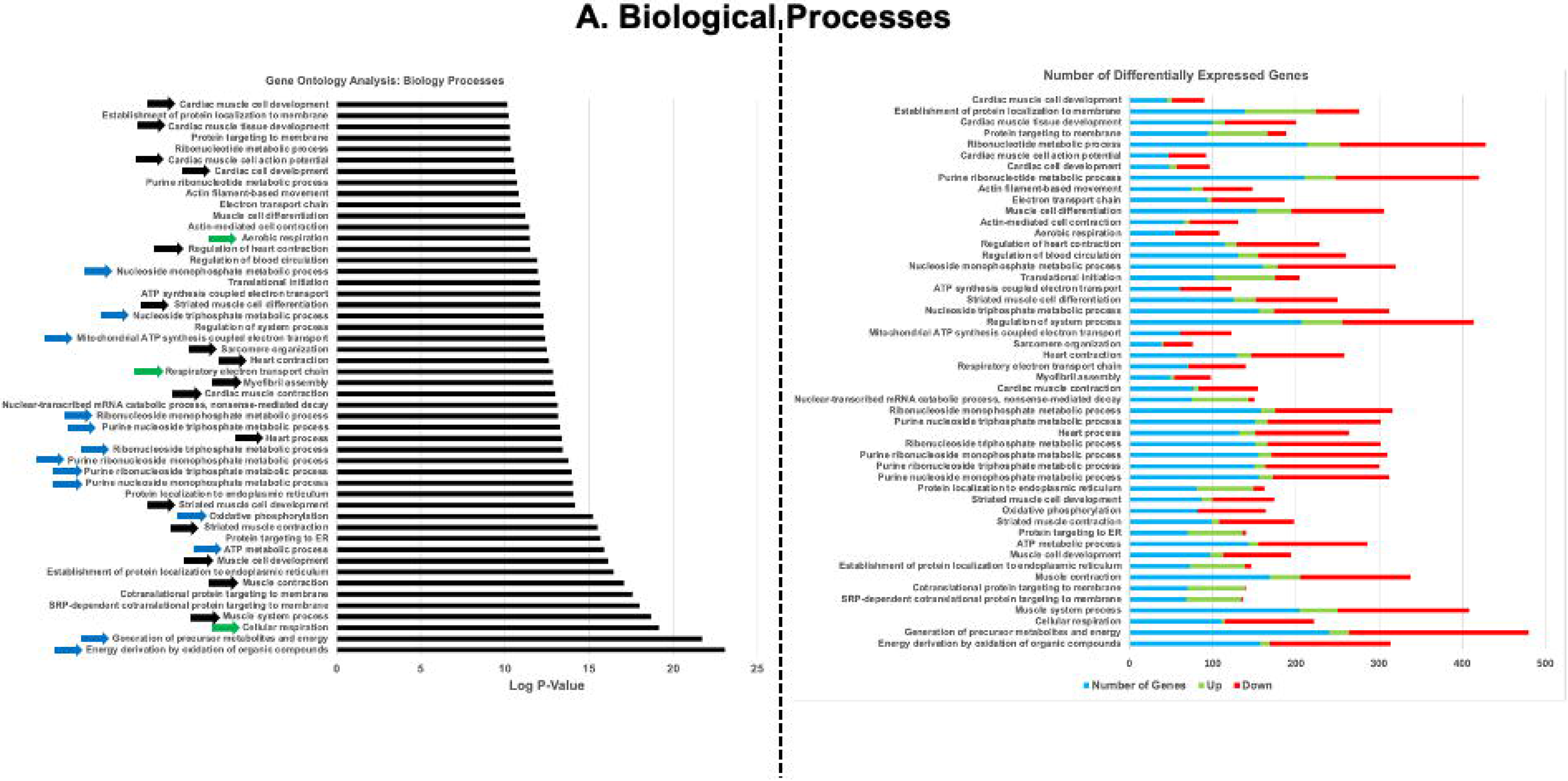
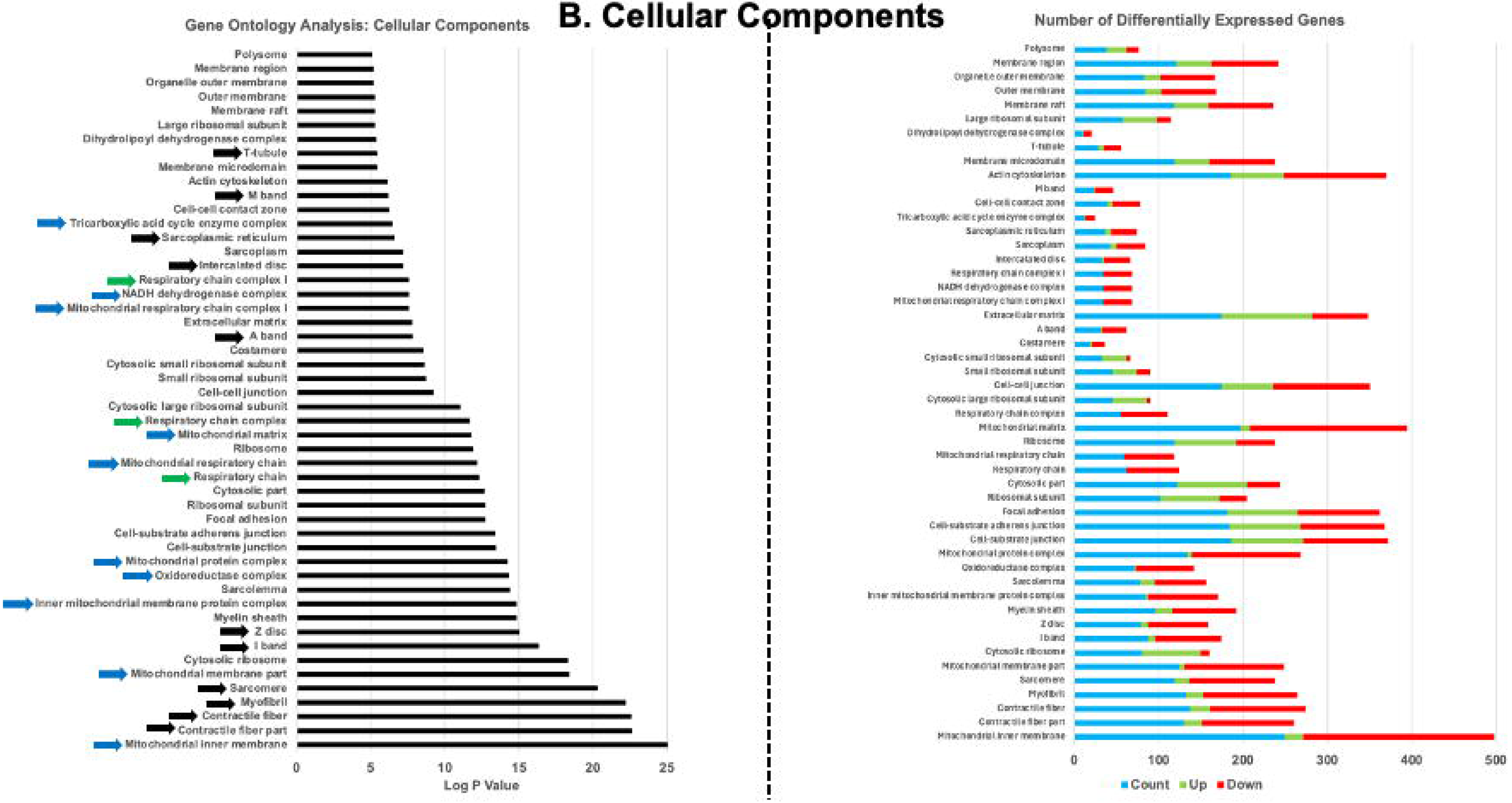
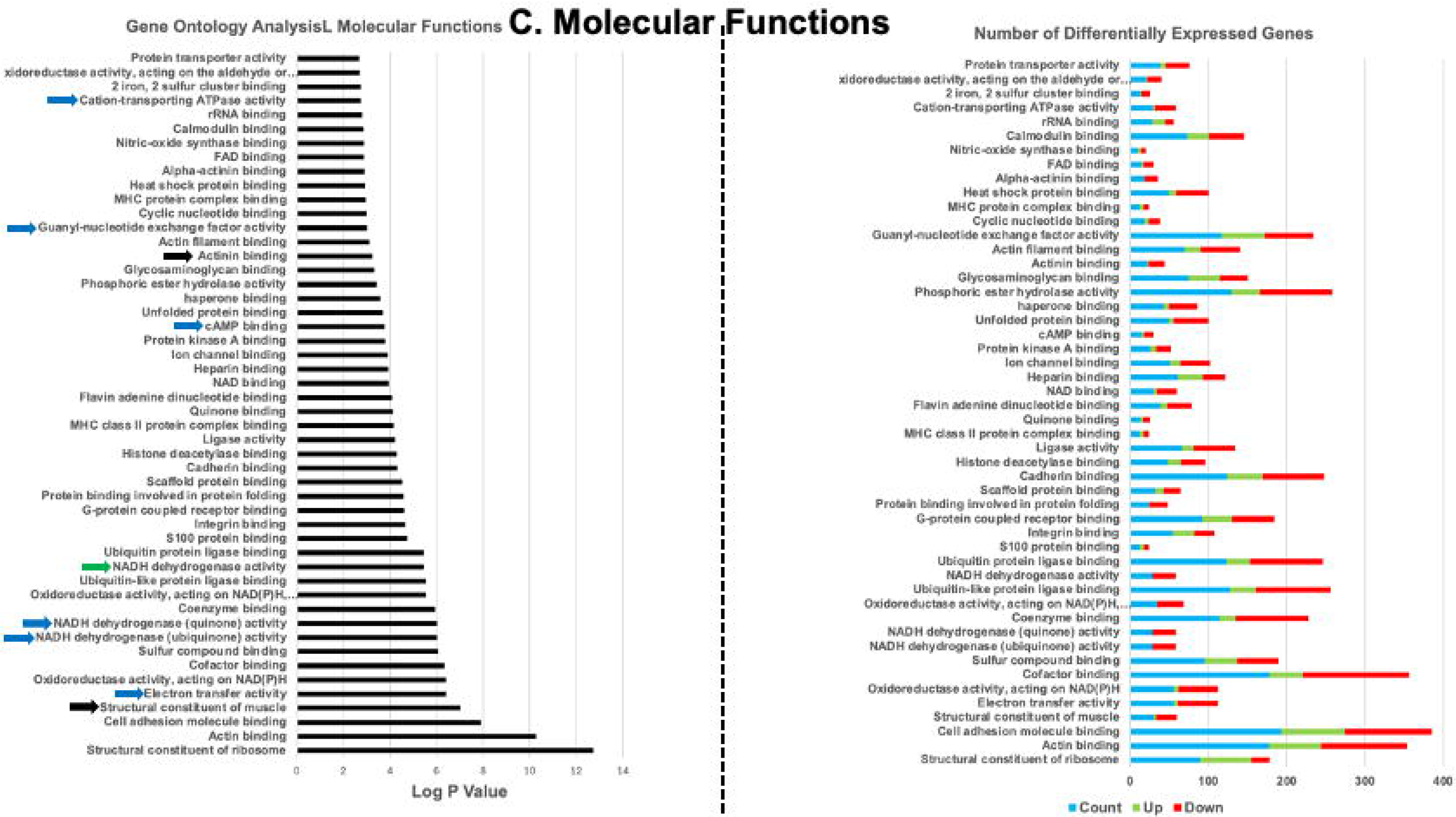
Gene Ontology Analysis for AggPPVS vs NonAggPPVS: (A) Biological Processes. (B) Cellular Components. (C) Molecular Functions. Figure on the left panel shows the top 50 most significant pathways. Figure on the right shows the number of differentially expressed genes for each pathway.

The GO analysis based on BPs showed upregulation of many pathways in AggPPVS relative to NonAggPPVS related to cardiac muscle contraction and function, respiration and energetics, all consistent with PVS phenotype. CCs components identified were mitochondria, cell membrane, extracellular matrix, cell-cell matrix, respiratory chain complex, mitochondrial matrix, ribosome, focal adhesion, mitochondrial protein complex, Z band, I band, sarcomere, myofibril and contractile fiber. Many of the these were consistent with changes in the contractile machinery of heart muscle tissue, energetics, respiratory and extracellular matrix remodeling. MFs were also consistent with BPs and CCs, including structural component of muscle, electron transfer activity, coenzyme binding, scaffold protein binding, cAMP binding, which as before, suggest changes in energetics, respiratory and heart muscle contraction.

### KEGG Enrichment Analysis

KEGG pathways are used to understand higher level function of cells and combined with GO analysis (as described earlier) and reactome pathways (described next), provides a comprehensive analysis of the changes in genomic profile changes related to PVS. For the KEGG enrichment analysis, the top 50 pathways, based on significance, were selected for discussion, and presented as a bar graph for AggPPVSvsNonAggPPVS (**Figure 6A**). The results from the KEGG analysis were consistent with the results from the GO analysis. Changes in energy metabolism, respiration and muscle contraction were notably different in group 1 (nonaggressive PVS) vs group 2 (aggressive PVS).

**Figure 6:**
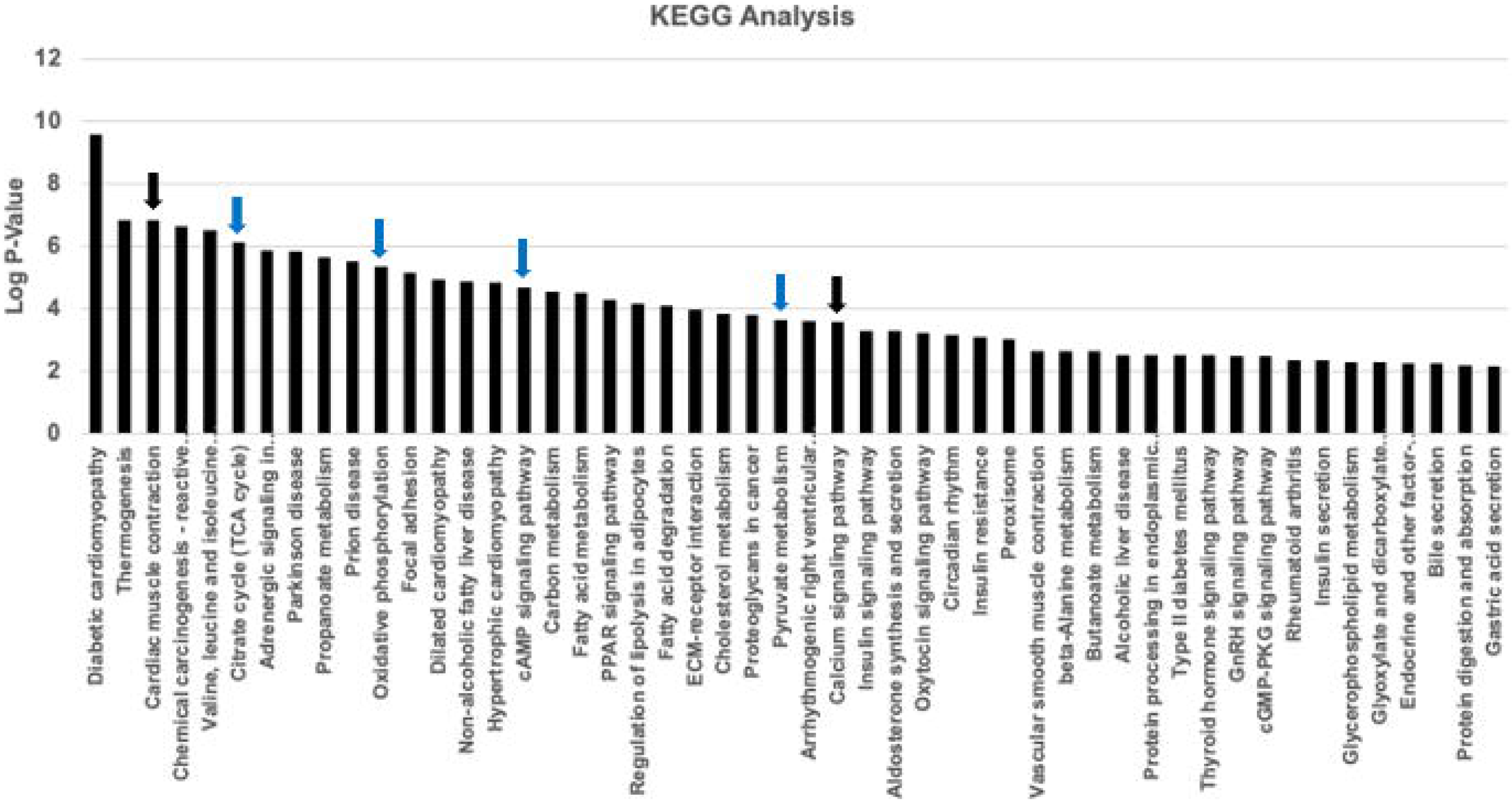
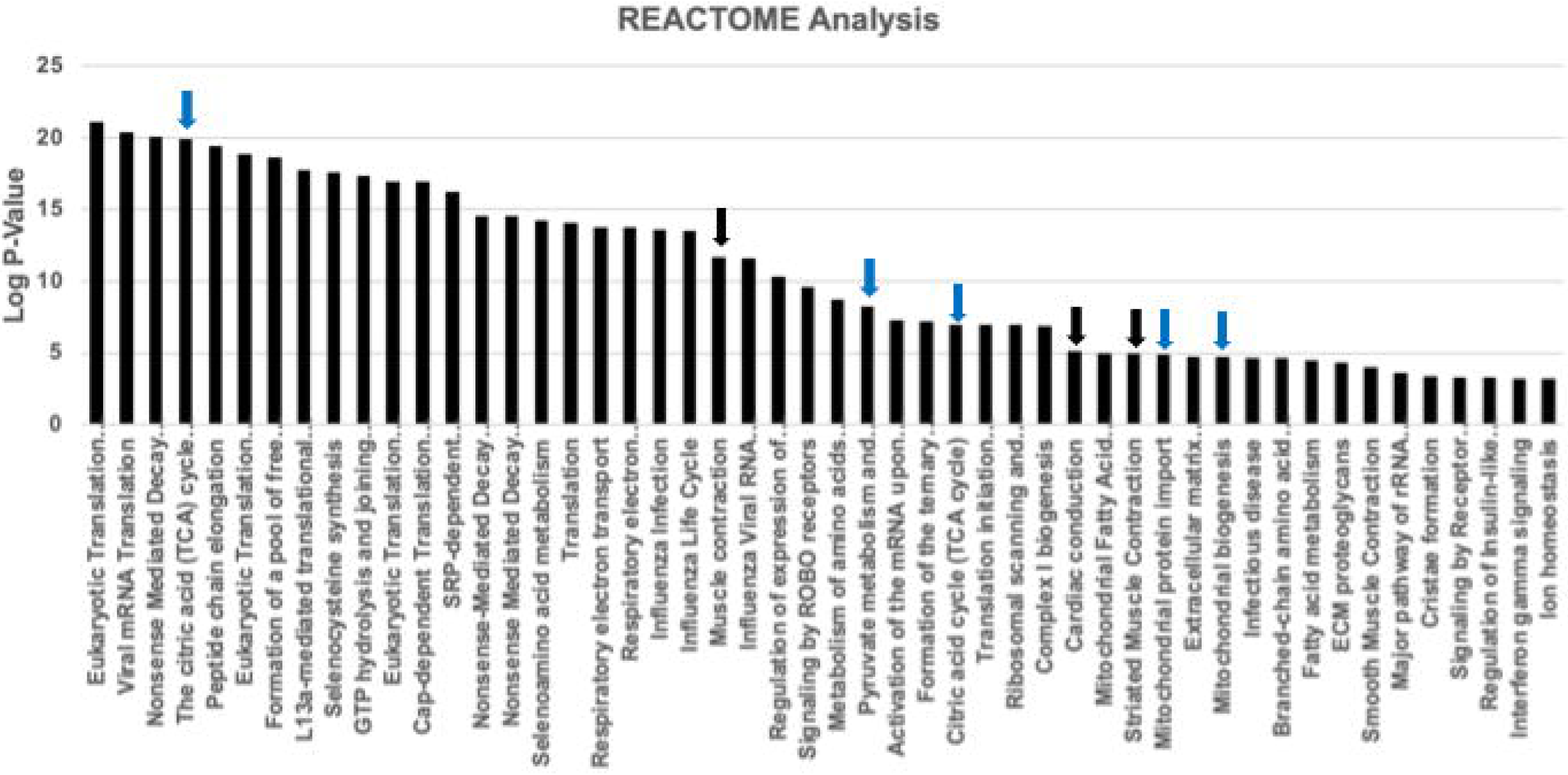
(A) KEGG Analysis for AggPPVS vs NonAggPPVS: (B) Reactome Analysis for AggPPVS vs NonAggPPVS. For both (A) and (B) the top 50 most significant pathways are shown.

### Reactome Pathway Analysis

To further validate the results from the GO analysis and the KEGG enrichment analysis, we conducted an additional test using the reactome pathway analysis, another powerful data mining tool to interpret bioinformatics data (**Figure 6B**). Not surprising, the reactome pathway analysis further validated our results, with energy metabolism, respiration and heart muscle contraction proving to be important when comparing AggPPVS vs NonAggPPVS.

### Selected Pathways

We selected 2 pathways and presented the specific gene that were either upregulated or downregulated (**Figure 7**). The specific pathways were selected were cardiac muscle contraction (**Figure 7A**) and calcium handling pathways (**Figure 7B**), selected to cover some of the important functions that change between AggPPVS vs NonAggPPVS, including heart muscle contraction and calcium handling.

**Figure 7:**
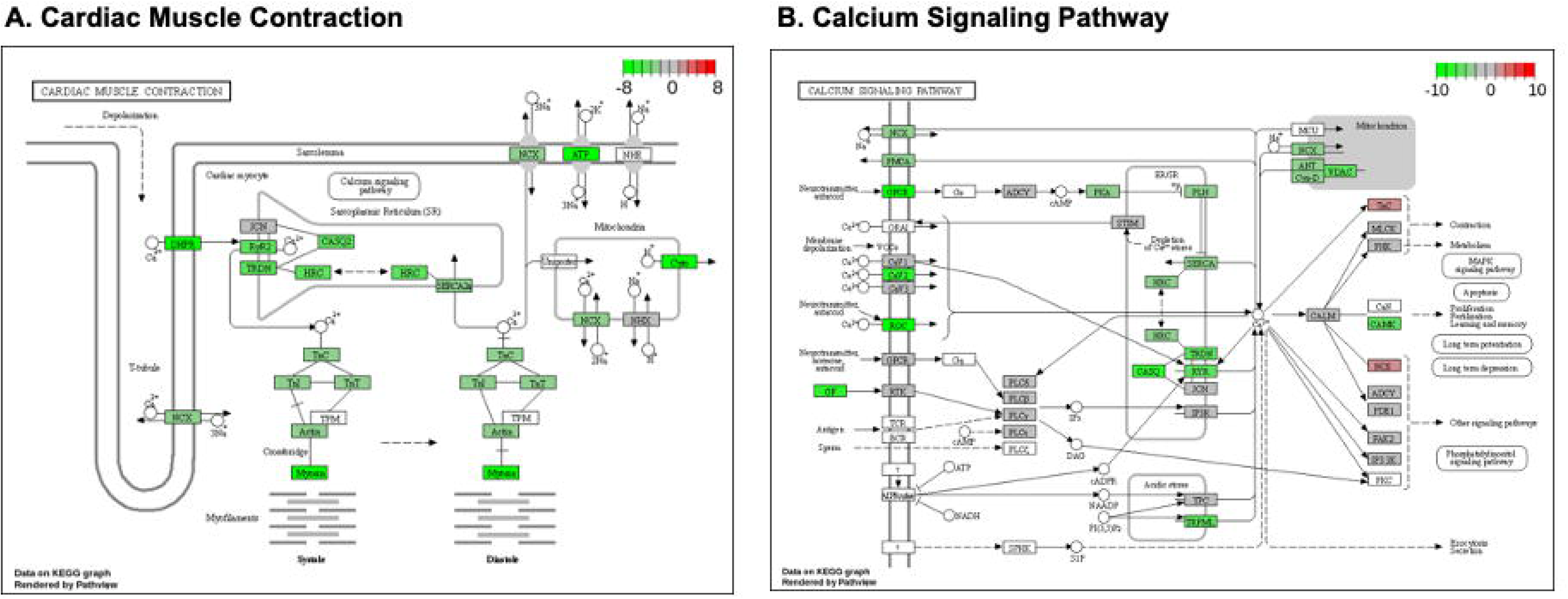
Selected KEGG Pathway for AggPPVSvsNonAggPPVS: (A) Cardiac Muscle Contraction. (B) Calcium Signaling Pathway.

## Discussion

PPVS and PRPVS are clinically distinct, with PPVS considered to be a congenital disorder and PRPVS acquired after surgical intervention. Both forms of PVS are difficult to manage and are associated with a high mortality rate in the pediatric population. Yet, the molecular basis of PPVS or PRPVS remain unknown. Changes in the expression of genes responsible for the onset of PPVS and PRPVS also remain unknown. The triggers that drive the differential expression of genes in PPVS and PRPVS are unknown. It is also not known if the same or a completely different set of genes are responsible for the initial onset of PPVS vs PRPVS. In addition, the genes responsible for the initial onset of the disease verses recurrent phenotype have not been identified. This study is attentive to these knowledge gaps in our understanding of PVS and is designed to identify differentially expressed genes responsible for both aggressive and nonaggressive PPVS and PRPVS.

There are many reasons to identify differentially expressed genes expression responsible for both aggressive and nonaggressive PPVS and PRPVS. First, such an understanding is critical to identifying the underlying the signaling pathways and subsequently develop new therapies to treat this patient population. Second, identification of specific markers can lead to the development of tools to predict the risk of recurrent phenotype and differentiation of management strategy based on risk profile. Third, identification of the genetic basis for PVS can lead to better tools for counseling for the families of affected patients.

Our first task was to identify differences and similarities between 8 comparison groups identified in **Figure 1F**. We made use of PCA analysis (**Figure 2**) and differential gene expression, visualized using volcano plots (**Figure 3**) to accomplish. Based on our analysis, we identified significant differences between four groups: AggPPVS vs NonAggPPVS, AllAgg vs AllNonAgg, AggPRPVS vs NonAggPRPVS, AggPPVS vs Ctrl. In addition, we identified similarities between four groups: AggPPVS vs AggPRPVS, NonAggPPVS vs NonAggPRPVS, NonAggPRPVS vs Ctrl and AllPPVS vs AllPRPVS. There were notable observations within this dataset. Clinically, AggPPVS and AggPRPVS are thought to be distinct, the former a congenital condition and the latter acquired after surgical intervention. However, our results show that AggPPVS and AggPRPVS are not distinct and may have overlapping genes and pathways responsible for mediating an aggressive phenotype. Similarly, NonAggPPVS vs NonAggPRPVS were also found to be similar, again suggesting common genes and pathways for nonaggressive primary and post-repair PVS. Another important finding was the similarity between AllPPVS vs AllPRPVS, suggesting common molecular triggers for both primary and post-repair PVS, irrespective of aggressive or nonaggressive phenotype.

Our initial analysis based on PCA analysis (**Figure 2**), and differential gene expression visualized using volcano plots (**Figure 3**) was used as a funneling strategy. We determined that four comparisons were significantly different and only these four were retained for further analysis. The four comparisons that were retained for further analysis were: AggPPVS vs NonAggPPVS, AllAgg vs AllNonAgg, AggPRPVS vs NonAggPRPVS, AggPPVS vs Ctrl. The next steps in our analysis were designed to identify specific genes and pathways in these four groups.

The results in **Figure 4A** show a significant upregulation of genes associated with collagen, including COL1A1, COL3A1, COL14A1 and COL1A2. This is consistent with the phenotype observed in PVS patients, with an abundance of ECM proteins identified within the fibrotic tissue in the pulmonary vein. Our results provide evidence that the fibrotic tissue consists of, at in in part, an increase in the expression of collagen.

We observed a significant increase in the expression of PIEZO1 in AggPPVS relative to NonAggPPVS. The log fold change in the expression of PIEZO1 was 2.19 or 4.55 in absolute terms (**Figure 4A**). In addition, PIEZO1 was one of the most significant genes based on p-value, determined to be 1.15×10^-12^. This is a very novel finding, as PIEZO1 has been linked to PVS for the first time through our study. Given that PIEZO1 is significantly overexpressed in aggressive PPVS patients relative to nonaggressive PPVS patients, this provides potential mechanistic insight. A lot is known about the activity of PIEZO1, and it is expressed in endothelial cells and acts as a mechanosensitive receptor^14^.

When exposed to elevated shear stress, activation of PIEZO1 results in an influx of intracellular calcium which results in changes in endothelial cell function^14^. Based on our data, PIEZO1 activation in PPVS patients appears to be involved in mediating an aggressive phenotype, and this hypothesis is further supported based on an increase in collagen production in this same patient population. While our data suggests this as a potential mechanism, additional validation studies will be required to validate this hypothesis. However, this is a very significant finding and provides preliminary evidence to support the role of PIEZO1 as a regulator of AggPPPVS and upon further validation, inhibition of the activity of PIEZO1 can potentially reduce the risk of recurrent phenotype in PPVS patients.

We next quantified DEGs in AllAgg vs AllNonAgg (**Figure 4B**). AggPRPVS vs NonAggPRPVS (**Figure 4C**) and AggPPVS vs Ctrl (**Figure 4D**). AllAgg and AllNonAgg contains samples from aggressive and nonaggressive primary and post-repair PVS, respectively. This comparison is designed to identify common genes responsible for recurrent phenotype, irrespective of disease origin (primary or post-repair). Genes associated with the production of collagen were upregulated and the same was true for PIEZO1, thereby suggesting that overexpression of PIEZO1 and subsequent ECM modeling through collagen production mediates aggressive phenotype in both primary and post-repair PVS. PIEZO1 was significantly upregulated in AggPRPVS relative to NonAggPRPVS with a log fold change of 2.34 and absolute fold change of 5.1. Since it was not one the top 50 significant genes, it was not a part of **Figure 4C**. Collectively, based on the DEGs for AggPPVS vs NonAggPPVS (**Figure 4A**), AllAgg vs AllNonAgg (**Figure 4B**) and AggPRPVS vs NonAggPRPVS (**Figure 4C**), we hypothesize that an increase in wall shear stress in the pulmonary vein of aggressive PVS patients, both primary and post-repair, results in an overexpression of PIEZO1 and subsequent production of collagen. Our final comparison in **Figure 4D** shows AggPPVS vs Ctrl and demonstrates a significant increase in genes associated with ECM production and remodeling, thereby providing further support for our hypothesis.

Once we identified differentially expressed genes, our next step in the analysis was to identify specific pathways that are responsible for mediating PVS phenotypes. Pathway analysis was only conducted on AggPPVS vs NonAggPPVS due to the clinical relevance; in a certain patient population, PPVS is observed independent of other conditions and a large percentage of these patients demonstrate recurrent phenotype. The GO analysis is presented in Figure 5. GO analysis consists of three components, BPs, which refers to the specific processes e.g. heart muscle contraction, CCs, the components of the cell responsible to mediate the specific BP, e.g., sarcomere and finally, MF, the specific molecular component responsible for the BPs e.g., electron transfer. The samples that were used for bulk-RNA sequencing were excised at the boundary between the pulmonary vein and left atrium and therefore, contained myocardium tissue in addition to pulmonary vein tissue. As is evident from **Figure 5A**, the vast majority of biological processes that were altered in in AggPPVS relative to NonAggPPVS were related to myocardial contractility and function, many pathways related to metabolism (blue arrows) and respiration (green arrows) were also significantly upregulated. The results of our GO analysis based on biological process demonstrate significant changes in pathways related to myocardial contraction, energy metabolism and respiration in AggPPVS relative to NonAggPPVS.

Our next step in the GO analysis was analysis of CCs (**Figure 5B**). These results demonstrate that many of the significantly upregulated cellular components in AggPPVS relative to NonAggPPVS were related to heart muscle contraction and consisted of the Z-disc, I-band, sarcomere, myofibril and contractile fiber (**Figure 5B**). Similar results were noted for changes in calcium handling with significant changes in t-tubule and sarcoplasmic reticulum in AggPPVS relative to NonAggPPVS (**Figure 5B**). In addition, results were also noted for CCs related to energy metabolism and respiration (**Figure 5B**). Collectively, the results of CCs analysis aligned well with the results of BPs (**Figure 5A**) and support the view the changes in heart muscle contractility, calcium transients, energy metabolism and respiratory are significantly difference in AggPPVS relative to NonAggPPVS. These results were also consistent with MPs analysis (**Figure 5C**) and provide further support for this hypothesis.

Our next step in the analysis was to identify KEGG (**Figure 6A**) and Reactome (**Figure 6B**) pathways that are significantly regulated in AggPPVS relative to NonAggPPVS. The results are consistent with the GO analysis and provide further evidence of the role of heart muscle contractility, calcium transients, energetics and metabolism processes in mediating AggPPVS relative to NonAggPPVS.

Given that the most significant pathways regulating AggPPVS relative to NonAggPPVS were cardiac contractility and intracellular calcium transients, specific differentially expressed genes in representative pathways were selected for further analysis (**Figure 7**). **Figure 7A** shows changes in specific genes in cardiac muscle contraction (**Figure 7A**), where myosin heavy chain expression is significantly downregulated during both systole and diastole. Given the known function of myosin in heart muscle contractility, the subsequent effect of this will be a decrease in the twitch force of contraction^15^. While myosin downregulation is the end result, many modulators of this are identified in **Figure 7A**, including actin, TnI, TnC and TnT, all of which are known to be key regulators of heart muscle contractility ^15^. Also shown **Figure 7A**, is a significant downregulation of proteins that regulate intracellular calcium, including RyR2 and CASQ2 and the same for membrane ion channels including the calcium pump DHPR and the calcium-sodium exchanger NCX. These results demonstrated downregulation in the expression of several key proteins known to be involved in cardiac contractility ^15^, many of which can serve as potential therapeutic targets.

Identifying DEGs during heart muscle contraction for AggPPVS relative to NonAggPPVS eluted to changes in intracellular calcium signaling (**Figure 7A**). We looked at the calcium signaling pathway (**Figure 7B**) to further quantify DEGs related to calcium signaling in cardiomyocytes. There was a significant downregulation in the expression of the calcium channel CaV2, the calcium uptake channel SERCA and CaMK, known to produce a cardioprotective effect on cardiomyocytes ^16, 17^ (**Figure 7B**). Collectively, these results further demonstrate the impaired calcium handling cascade in heart muscle tissue from AggPPVS samples relative to NonAggPPVS.

Collectively, our results lead us to postulate a potential mechanism for PVS. Our proposed mechanism applies for both PPVS and PRPVS and our data suggests a common mechanism for both phenotypes. Furthermore, our work identified a potential mechanism to explain aggressive PPVS and PRPVS relative to nonaggressive PPVS and PRPVS. Our work has identified PIEZO1 as a potential mediator of aggressive PPVS and PRPVS and though a trigger has not been identified, we speculate elevated shear stress to be the mediator of PIEZO1 overexpression, based on the known function of this transmembrane protein as a mechanosensitive receptor. It is also possible, and highly likely that PIEZO1 is only one mediator of aggressive PPVS and PRPVS and other mediators will play a role. The effector of our proposed mechanism has been identified as diminished contractile function and reduced expression of genes responsible for intracellular calcium regulation. This explains the diminished heart muscle function observed during PVS.

There are limitations to the current study. While PIEZO1 expression was significantly upregulated in aggressive PPVS and PRPVS tissue relative to nonaggressive PPVS and PRPVS tissue, the role of PIEZO1 has not been validated through other techniques like single cell sequencing and downstream regulators not identified. Given the rarity of this disease process and introduction of sutureless repair techniques, sample sizes and lack of control pulmonary vein tissue were a substantial limitation in study design. Multicenter collaboration is necessary to increase access to patient samples and increase our understanding of the molecular mechanisms responsible for PVS. Recent advances in stem cell technology ^18–20^, organoid culture^21, 22^ and bioengineered heart muscle ^23–25^ need to be leveraged to provide additional mechanistic insight. Bioreactors need to be developed that accurately replicate the complex fluid shear stresses within the pulmonary vein of PVS patients ^26–28^.

### Conclusions

The results of this study provide insight into the potential mechanism of aggressive PPVS and PRPVS relative to nonaggressive PPVS and PRPVS. Based on our data, an increase in wall shear stress causes an upregulation in the expression of PIEZO1, which triggers an intracellular signaling cascade that results in reduced contractility and calcium handling of heart muscle tissue.

## Acknowledgements

The authors would like to acknowledgement financial support from the Division of Congenital Heart Surgery for the completion of this study.

## Glossary of abbreviations

PAPVC: partial anomalous pulmonary venous connection
PR-PVS: post-repair pulmonary vein stenosis
PVS: pulmonary vein stenosis
PPVS: primary pulmonary vein stenosis
RNA-seq: bulk radionucleic acid sequencing
TAPVC: total anomalous pulmonary venous connection
EndoMT: endothelial to mesenchymal transition
ECM: extracellular matrix
GO: gene ontology
BP: biological processes
CC: cellular components
MF: molecular function

## Notes

### Competing Interest Statement

The authors have declared no competing interest.

